# Systematic visualisation of molecular QTLs reveals variant mechanisms at GWAS loci

**DOI:** 10.1101/2023.04.06.535816

**Authors:** Nurlan Kerimov, Ralf Tambets, James D. Hayhurst, Ida Rahu, Peep Kolberg, Uku Raudvere, Ivan Kuzmin, Anshika Chowdhary, Andreas Vija, Hans J. Teras, Masahiro Kanai, Jacob Ulirsch, Mina Ryten, John Hardy, Sebastian Guelfi, Daniah Trabzuni, Sarah Kim-Hellmuth, Will Rayner, Hilary Finucane, Hedi Peterson, Abayomi Mosaku, Helen Parkinson, Kaur Alasoo

## Abstract

Splicing quantitative trait loci (QTLs) have been implicated as a common mechanism underlying complex trait associations. However, utilising splicing QTLs in target discovery and prioritisation has been challenging due to extensive data normalisation which often renders the direction of the genetic effect as well as its magnitude difficult to interpret. This is further complicated by the fact that strong expression QTLs often manifest as weak splicing QTLs and vice versa, making it difficult to uniquely identify the underlying molecular mechanism at each locus. We find that these ambiguities can be mitigated by visualising the association between the genotype and average RNA sequencing read coverage in the region. Here, we generate these QTL coverage plots for 1.7 million molecular QTL associations in the eQTL Catalogue identified with five quantification methods. We illustrate the utility of these QTL coverage plots by performing colocalisation between vitamin D levels in the UK Biobank and all molecular QTLs in the eQTL Catalogue. We find that while visually confirmed splicing QTLs explain just 6/53 of the colocalising signals, they are significantly less pleiotropic than eQTLs and identify a prioritised causal gene in 4/6 cases. All our association summary statistics and QTL coverage plots are freely available at https://www.ebi.ac.uk/eqtl/.

## Introduction

Most genetic variants associated with complex traits are in the non-coding regions of the genome (Maurano et al., 2012). More than a decade of molecular quantitative trait locus (QTL) studies has revealed that these variants regulate either the expression level (Kerimov et al., 2021; The GTEx Consortium, 2020), splicing (Li et al., 2016), promoter usage (Alasoo et al., 2019; Garieri et al., 2017) or alternative polyadenylation (Mittleman et al., 2020; Yoon et al., 2012) of their target genes. Although the eQTL Catalogue has contained transcript-level QTL summary statistics from the beginning, characterising the exact mechanism of action of each molecular QTL has remained challenging due to considerable overlap between QTLs detected by different RNA-seq quantification methods (Kerimov et al., 2021), technical biases in read alignment (van de Geijn et al., 2015), and a large number of alternative transcripts or splice junctions to be considered for each gene. Furthermore, because the usage of each transcript or splice junction is quantified relative to all other transcripts, the magnitude and direction of the genetic effect, the part of the gene affected, as well as the absolute expression of the affected transcript is often difficult to assess from summary statistics alone.

This ambiguity can be reduced by visualising the change in the average RNA-seq read coverage in the gene region associated with each additional copy of the alternative allele. We and other have used these QTL coverage plots to characterise chromatin QTLs (Alasoo et al., 2018; Degner et al., 2012; Kumasaka et al., 2018) as well as to confirm promoter usage and splicing QTLs (Alasoo et al., 2019). However, previous studies have visualised individual molecular QTLs in a setting where access to individual-level genotype and read coverage data is available. It has not been done systematically in large molecular QTL compendia such as the GTEx project (The GTEx Consortium, 2020) and the eQTL Catalogue, because in a naive implementation the read coverage stratification by genotype needs to be performed separately for each significant genetic variant and molecular trait pair of interest. Since transcript and exon-level analyses profile hundreds of thousands of correlated molecular traits in a single dataset, this means that the number of QTL coverage plots required can quickly become intractable.

In this update to the eQTL Catalogue, we present an approach to generate QTL coverage plots for all independent genetic signals and their associated molecular traits. First, we have updated our data processing workflows to improve promoter usage and splicing QTL discovery and to generate read coverage signals for all 25,724 RNA-seq samples. We have also adopted fine-mapping-based filtering to identify all independent genetic signals and associated molecular traits for each gene while reducing the size of the summary statistics files by 98%. Finally, to support new colocalisation methods that can account for multiple independent causal variants (Wallace, 2021), we have computed signal-level log Bayes factors for all independent signals (Wang et al., 2020). This approach has enabled us to predefine tag variants for all independent genetic associations identified in 127 eQTL datasets and generate QTL coverage plots that can be used to interpret almost all colocalising signals detected in the eQTL Catalogue.

## Results

### Updates to the eQTL Catalogue resource

The aim of the eQTL Catalogue is to provide a public resource of uniformly processed molecular QTL summary statistics and continuously update this resource as new studies, reference annotations and quantification methods become available. Here, we present the updates to the eQTL Catalogue release 6 that we have made since the publication of the original paper (release 3).

#### Newly added datasets

We have added nine new RNA-seq studies and one microarray study to the eQTL Catalogue. This has increased the total number of studies in the resource to 31, the total number of datasets to 127 and the cumulative number of donors and samples to 7,526 and 30,602, respectively (Figure 1A). Newly added datasets include additional datasets from tissues and cell types already present in the eQTL Catalogue (e.g. various brain regions (Guelfi et al., 2020; Hoffman et al., 2019), immune cells (Bossini-Castillo et al., 2022; Gilchrist et al., 2022; Kim-Hellmuth et al., 2017; Theusch et al., 2020) and induced pluripotent stem cells (DeBoever et al., 2017; Pashos et al., 2017)) as well as previously missing microglia (Young et al., 2021), placenta (Peng et al., 2018), hepatocytes (Pashos et al., 2017), and cartilage and synovium tissues (Steinberg et al., 2021). Complete summary of the datasets present in the eQTL Catalogue is shown in Supplementary Table 1.

**Figure 1.**
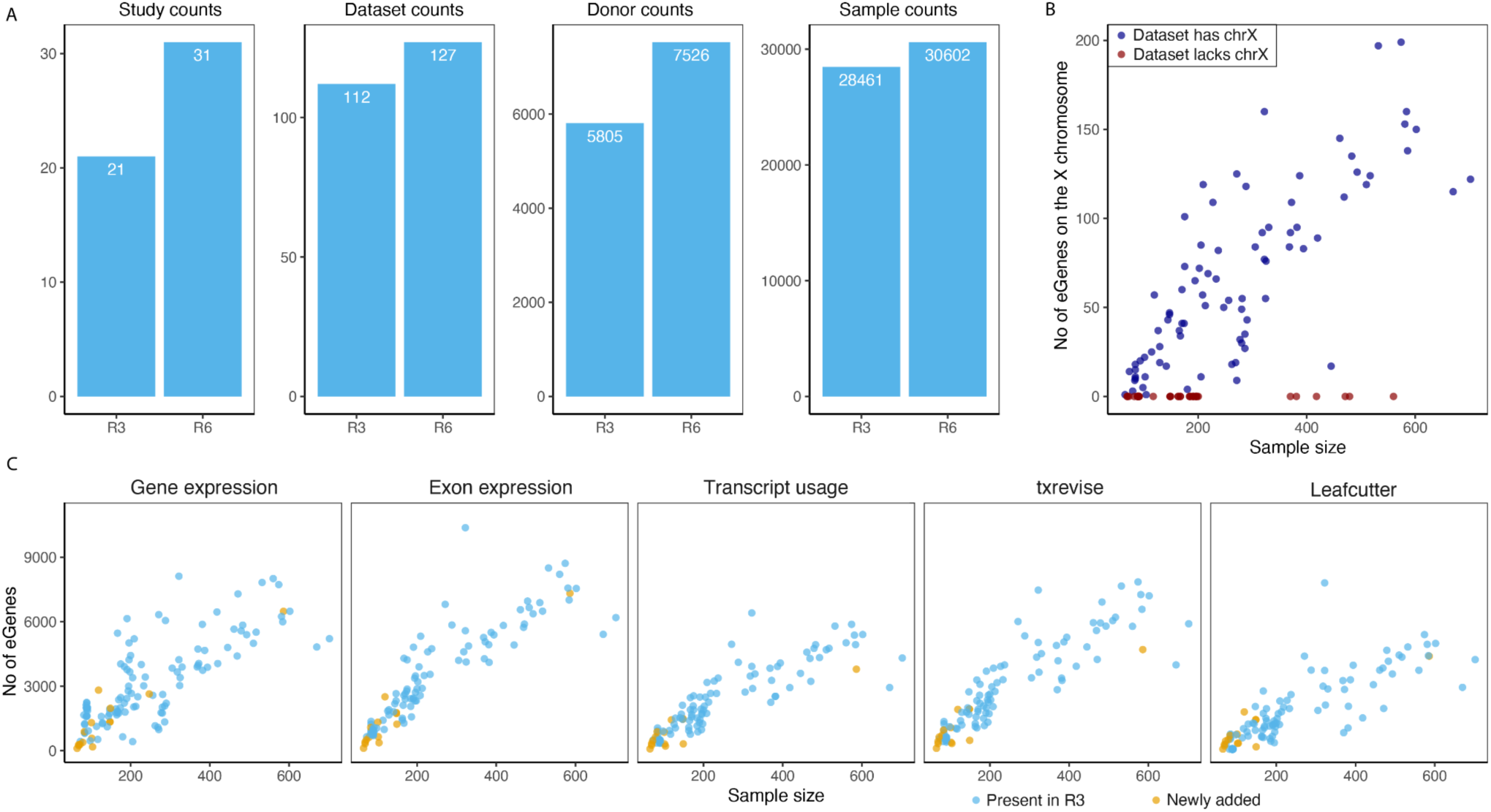
Uniform re-processing of all datasets. (**A**), The number of studies, datasets, donors, and samples in the previous publication (R3) and current version of the eQTL Catalogue (R6). **(B**), Number of genes with at least one significant eQTL (‘eGenes’) on the X chromosome as a function of dataset sample size. Red points indicate datasets for which the X chromosome genotypes were unavailable. (**C),** The number of eGenes identified in each dataset for the five molecular traits (gene expression, exon expression, transcript usage, txrevise event usage, and Leafcutter splice-junction usage). Datasets newly added since release 3 have been highlighted.

#### Imputation of the X chromosome genotypes

In addition to integrating new datasets, we also made two major changes to our genotype imputation workflow. First, we migrated to the new 1000 Genomes 30x of GRCh38 reference panel (Byrska-Bishop et al., 2022). This allowed us to impute genotypes directly to the GRCh38 build and avoid errors caused by the genomic coordinate lift over process. Secondly, our imputation workflow now also supports the X chromosome. As a result, summary statistics for 18 of the 31 studies now also contain variants from the X chromosome. Across these 18 datasets, we detected at least one significant eQTL (FDR <1%) for 853 unique genes on the X chromosome (Figure 1B). These X chromosome eQTLs account for ∼1.6% of all significant eQTLs (FDR <= 1%). Ten of the other 13 studies are missing the X chromosome QTLs because X chromosome genotypes were not deposited with data. Exceptions are male only studies (n = 3) that did not pass our genotype QC criteria (Supplementary Table 2).

#### Improved quantification of splicing and promoter usage QTLs

The previous release of the eQTL Catalogue included four molecular trait quantification methods to measure transcriptional changes from RNA-seq data: gene expression (ge), exon expression (exon), transcript usage (tx) and transcriptional event usage (txrevise). In addition to these four, we have now also implemented LeafCutter (Li et al., 2018) to directly quantify the usage of splice junctions (Supplementary Figure 1). We have also augmented the txrevise promoter annotations with experimentally determined promoters from the FANTOM5 project (Vija and Alasoo, 2022). Finally, we have updated the reference transcriptome annotations to Ensembl version 105 and GENCODE version 39. We observed a clear linear relationship between the number of significant associations detected with each quantification method and the dataset sample size, with gene expression, exon expression and txrevise detecting, on average, slightly more associations than transcript usage and Leafcutter (Figure 1C).

#### Fine-mapping-based filtering of transcript-level summary statistics

A major challenge in working with exon- and transcript-level (transcript usage, txrevise, leafcutter) associations is the large number of correlated traits being tested that result in very large summary statistics files. For example, typical summary statistics for exon and txrevise QTLs are 15-20 times larger than the corresponding files for gene expression QTLs. In addition to complicating our data release and archival procedures, these large file sizes meant that performing comprehensive colocalisation analysis against the eQTL Catalogue required the downloading and processing of >15Tb of data. To reduce the size of these files, we have now implemented fine-mapping-based filtering. Briefly, we are using fine mapped credible sets to identify all independent signals at the gene level. We then filter the summary statistics files to only retain the most strongly associated molecular trait (exon, transcript, txrevise event or Leafcutter splice junction) for each signal. This filtering reduces the size of the summary statistics files for those quantification methods by ∼98% while retaining almost all significant associations for colocalisation purposes. Reducing the size of the univariate summary statistics files has also allowed us to export SuSiE log Bayes factors for each fine mapped signal and all tested variants (Wang et al., 2020). As illustrated below, these log Bayes factors can be directly used in the new coloc.susie method to perform colocalisation analysis between all pairs of independent signals (Wallace, 2021).

#### Visualisation of transcript-level associations

Another benefit of fine-mapping-based filtering is that we now have a tractable set of independent lead variants and associated molecular traits across all datasets and quantification methods that we can visualise using static QTL coverage plots. These plots display normalised RNA-seq read coverage across all exons of the gene (Figure 2A), exon-level QTL effect sizes and standard errors (Figure 2B), as well as the alternative transcripts or splice junctions used in association testing (Figure 2C). As an example, we are highlighting the association between chr11_14855172_G_A and alternative splicing of exon four of the *CYP2R1* gene (Figure 2). The static QTL coverage plots for all 1,716,482 independent signals are now available via the eQTL Catalogue FTP server.

**Figure 2.**
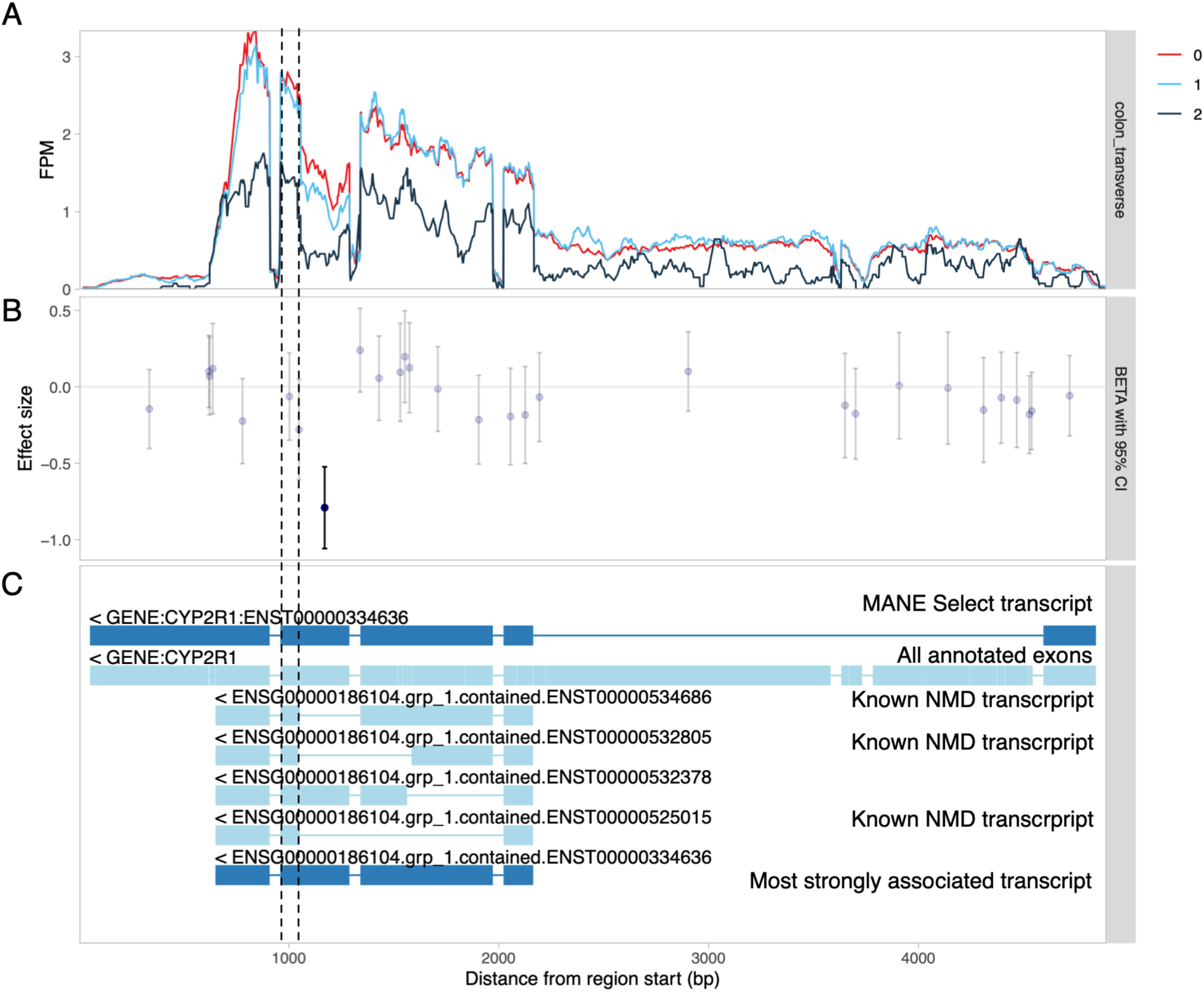
Visualisation of a splicing QTL detected in the *CYP2R1* gene. (**A**) RNA-seq read coverage across the *CYP2R1* gene in GTEx transverse colon tissue stratified by the genotype of the lead sQTL variant (chr11_14855172_G_A). All introns have been shortened to 50 nt with wiggleplotr (Alasoo, 2017) to make variation in exonic read coverage easier to see. (**B**) Effect sizes and 95% confidence intervals of the lead sQTL variant on the expression level of individual exons (or exonic parts) of *CYP2R1*. Associations significant at FDR <= 1% are shown in dark blue. (**C**) The top two rows show the MANE Select (Morales et al., 2022) reference transcript and all annotated exons of *CYP2R1*, respectively. The remaining rows show the txrevise (Alasoo et al., 2019) event annotations used for sQTL mapping. The short version of exon 4 (between dashed lines) is only present in annotated nonsense-mediated decay (NMD) transcripts.

### Case study: target gene prioritisation for vitamin D GWAS

To test the utility of the new QTL coverage plots, we performed a proof-of-concept colocalisation analysis between all molecular traits in the eQTL Catalogue and vitamin D levels in the UK Biobank. We chose this phenotype, because the vitamin D biosynthesis pathway is well understood and many causal genes underlying GWAS associations for vitamin D are already known (Hyppönen et al., 2022; Manousaki et al., 2020). At a stringent threshold of PP4 > 0.9, we found that 53/83 signals from 34/48 regions colocalised with 81 protein coding genes (Figure 3A). Although colocalisation with total gene expression was most common, there was considerable overlap between colocalisations detected with the five different quantification methods (Figure 3A).

**Figure 3.**
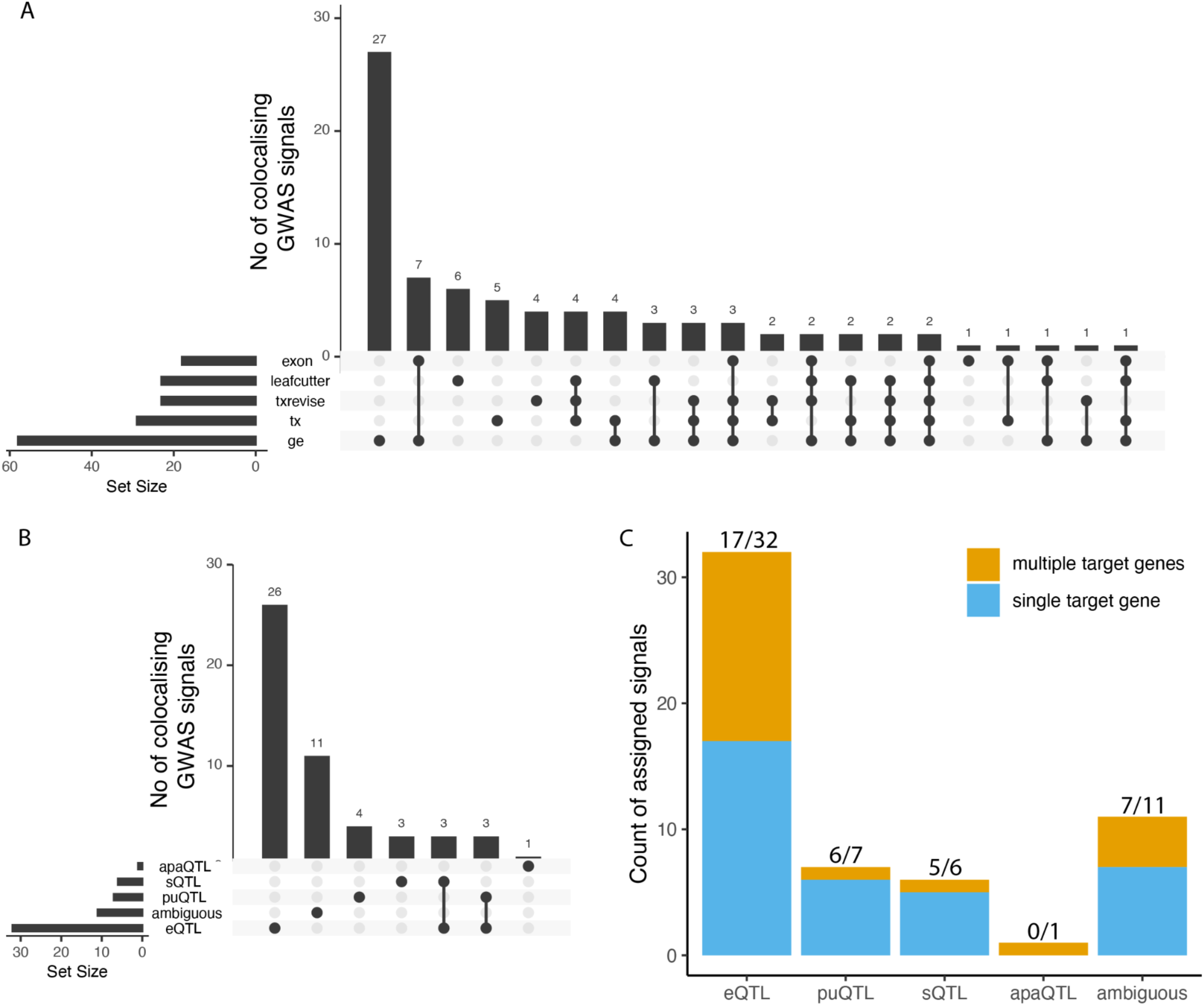
Sharing of significantly colocalised signals with vitamin D (**A**) Number of colocalised signals detected by the different molecular QTL quantification methods and sharing between them. (**B**) Number of colocalised signals assigned to empirical functional consequence (eQTL, sQTL, puQTL, apaQTL or ambiguous) and sharing structure between them. (**C**) Number of independent colocalised signals associated with either a single target gene or multiple target genes in each functional consequences group. eQTL - expression QTL, sQTL - splicing QTL, puQTL - promoter usage QTL, apaQTL - alternative polyadenylation QTL.

We extracted QTL coverage plots for all 816 colocalising molecular QTL signals. We then manually reviewed the plots to classify each signal into one of five categories: expression QTLs, promoter usage QTLs (puQTLs), splicing QTLs (sQTLs), alternative polyadenylation QTLs (apaQTLs) and ambiguous (Figure 3B). As an example, we detected a splicing QTL affecting the length of exon 4 of *CYP2R1* (Figure 2). *CYP2R1* is highly likely to be the causal gene at this locus as it codes the Cytochrome P450 2R1 microsomal vitamin D 25-hydroxylase (Cheng et al., 2003). We found that although transcript-level methods (tx, txrevise and leafcutter) detected at least one colocalisation for 37/53 independent signals, only 14 of those (7 puQTLs, 6 sQTLs and 1 apaQTL) could be classified as primary transcript-level QTLs (Figure 3B). Other 23 cases were either ambiguous or could be better explained by strong primary eQTL effects that led to small downstream changes in splicing or transcript usage (Supplementary Table 3).

Even though Leafcutter detected all seven visually confirmed sQTLs and 5/7 puQTLs, it also detected 11 additional signals, nine of which would be better explained by a strong eQTL effects (e.g. *CELSR2* eQTL at the *SORT1* locus (Supplementary Figure 2)). Thus, the fact that colocalisation is detected by one of the transcript-level methods (tx, txrevise or leafcutter) does not reliably indicate that the underlying signal is driven by a splicing mechanism. The visualisations also helped us to detect three likely cases of reference mapping bias at the *DHCR7*, *NUDT9* and *JUND* genes (Supplementary Figures 3-5). For discussion of why we opted not to correct for reference mapping bias during molecular trait quantification, see Supplementary Note.

We also noticed that 15/32 confirmed eQTL colocalised with more than one gene (Figure 3C). In contrast, only one of seven puQTLs and one of six sQTLs colocalised with multiple genes, suggesting that sQTLs and puQTLs might be less pleiotropic than eQTLs. To evaluate if lower pleiotropy also translated into more accurate causal gene prioritisation, we manually reviewed all of the 53 GWAS signals to identify the most likely causal genes. We integrated information about missense variant associations, gene presence in the vitamin D synthesis pathway and other literature evidence to prioritise the most likely causal gene for 28/53 GWAS signals (Supplementary Table 3). For four of the six sQTL signals, the colocalising gene overlapped the prioritised causal gene (*CYP2R1*, *HAL*, *GC* and *SDR42E1)* and for two signals we could not prioritise the causal gene. For eQTLs, we prioritise the most likely causal gene at 19/32 loci. In 11/19 cases (3 shared with sQTLs, Figure 3B) the colocalising eQTL genes completely overlapped the prioritised genes. In four cases the prioritised gene (*SORT1*, *FLG*, *HAL*, *CETP*) was one of multiple co-localizing genes. Finally, at four additional signals, the prioritised gene was different from the one that had eQTL colocalisation evidence (Supplementary Table 3). Interestingly, in three of the four cases the GWAS lead variant was a missense (*SEC23A*, *PLA2G3*) or a synonymous variant (*CYP2R1)* in the prioritised gene. While the number of loci observed here is small, these results suggest that while visually confirmed sQTLs colocalise with a smaller fraction of GWAS loci than eQTLs (6 vs 32), they are also less pleiotropic and thus more likely to identify the correct causal gene.

## Discussion

We have made three major changes to the eQTL Catalogue in release 6. First, we have integrated data from ten additional eQTL studies bringing the total number of unique eQTL datasets to 127. These datasets contain uniformly processed results from 30,602 samples from 7,526 individuals. We have also updated our genotype imputation, RNA-seq analysis and QTL mapping workflows to add support for the X chromosome, added Leafcutter as a splicing quantification method and added support for fine mapping-based colocalisation analysis with coloc.susie (Wallace, 2021). Finally, we have developed static QTL coverage plots to visualise molecular QTL associations at the level of RNA-seq read alignments. All our results and data are available on the eQTL Catalogue FTP server and REST API.

To quantify the impact of these updates, we performed colocalisation between all molecular QTLs present in the eQTL Catalogue and fine mapped GWAS signals for plasma vitamin D levels in the UK Biobank (Kanai et al., 2021). The QTL coverage plots allowed us to assign an empirical functional consequence (eQTL, sQTL, puQTL, apaQTL) for 42/53 colocalising loci while 11 remained ambiguous. This revealed that while primary sQTLs explained fewer GWAS signals than eQTLs, they also appeared to be less pleiotropic and more likely to identify the correct target genes. A limitation of our approach is that we used manual visual inspection to assign mechanisms to different types of molecular QTLs. Although we tried to be careful, there is a small risk that this approach could have introduced inadvertent confirmation bias (e.g. classifying less pleiotropic loci as sQTLs). We expect that it might be possible to automate this classification in the future by machine learning approaches that consider variant-level annotations such as splicing scores (Jaganathan et al., 2019; Zeng and Li, 2022) or distance to genomic features.

We also observed that while most GWAS signals colocalised with an eQTL, approximately ∼50% of the eQTL colocalisations prioritised more than one gene. Similarly in 4/19 cases, the colocalising gene was different from the manually prioritised causal gene. This agrees with multiple previous observations that eQTL colocalisation alone often achieves low precision in causal gene identification (Mountjoy et al., 2021; Nasser et al., 2021). This does not seem to be a simple artefact of colocalisation analysis as CRISPR experiments have also revealed that targeting a single enhancer often regulates the expression of multiple target genes (Engreitz et al., 2016; Fulco et al., 2019; Kasela et al., 2021). We believe that while eQTL colocalisation can sometimes reveal trait-relevant tissues or cell types, target gene identification requires integration of multiple strands of evidence. Considering variants with potentially less pleiotropic effects such as missense and splice variants can also be helpful.

The systematic re-analysis and visualisation of molecular QTLs presented here would not have been possible without the researchers of the 31 original studies making their individual-level gene expression and genotype data available for qualified researchers. We are committed to sharing all summary statistics and fine mapping results openly and will seek to continuously integrate new eQTL datasets as they become available. We are also working on making the static QTL coverage plots available via an API and an interactive web interface.

## Methods

### Data access and informed consent

For all newly added datasets, we applied for access via the relevant Data Access Committees. The database accessions and contact details of the individual Data Access Committees can be found on the eQTL Catalogue website (http://www.ebi.ac.uk/eqtl/Studies/). In our applications, we explained the project and our intent to share the association summary statistics publicly. Ethical approval for the project was obtained from the Research Ethics Committee of the University of Tartu (approval 287/T-14).

### Genotype data

#### Pre-imputation quality control

We lifted coordinates of the genotyped variants to the GRCh38 build with CrossMap v0.4.1 (Zhao et al., 2014). We aligned the strands of the genotyped variants to the 1000 Genomes 30x on GRCh38 reference panel (Byrska-Bishop et al., 2022) using Genotype Harmonizer (Deelen et al., 2014). We excluded genetic variants with Hardy-Weinberg p-value < 10^-6^, missingness > 0.05 and minor allele frequency < 0.01 from further analysis. On the X chromosome, we applied the QC filters to female samples only and then retained the same variants also in the male samples. We also excluded samples with more than 5% of their genotypes missing.

#### Genotype imputation and quality control

We pre-phased and imputed the microarray genotypes to the 1000 Genomes 30x on GRCh38 reference panel (Byrska-Bishop et al., 2022) using Eagle v2.4.1 (Loh et al., 2016) and Minimac4 (Das et al., 2016). On the X chromosome, we performed imputation separately for variants located in the pseudoautosomal (PAR) and non-PAR regions. After imputation, we multiplied male genotype dosage in the non-PAR region by two to ensure that it is on the same scale with the female genotypes. We used bcftools v1.9.0 to exclude variants with minor allele frequency (MAF) < 0.01 and imputation quality score R2 < 0.4 from downstream analysis. The genotype imputation and quality control steps are implemented in <u>eQTL-Catalogue/genimpute</u> (v22.01.1) workflow available from GitHub.

We aligned the low-coverage whole genome sequencing (WGS) data from the BLUEPRINT project to the GRCh38 reference genome with bwa v0.7.17 (Li, 2013) and performed imputation to the 1000 Genomes 30x on GRCh38 reference panel using GLIMPSE v1.1.1 (Rubinacci et al., 2021). The low-coverage WGS genotype imputation workflow is available from GitHub: https://github.com/peepkolberg/glimpse.

### Phenotype data

#### Studies

eQTL Catalogue release 6 contains phenotype data from the following 25 RNA-seq: ROSMAP (Ng et al., 2017), BrainSeq (Jaffe et al., 2018), TwinsUK (Buil et al., 2015), FUSION (Taylor et al., 2019), BLUEPRINT (Chen et al., 2016; Kundu et al., 2020), Quach_2016 (Quach et al., 2016), Schmiedel_2018 (Schmiedel et al., 2018), GENCORD (Gutierrez-Arcelus et al., 2013), GEUVADIS (Lappalainen et al., 2013), Alasoo_2018 (Alasoo et al., 2018), Nedelec_2016 (Nédélec et al., 2016), Lepik_2017 (Lepik et al., 2017), HipSci (Kilpinen et al., 2017), van_de_Bunt_2015 (van de Bunt et al., 2015), Schwartzentruber_2018 (Schwartzentruber et al., 2018), GTEx v8 (The GTEx Consortium, 2020), CAP (Theusch et al., 2020), Peng_2018 (Peng et al., 2018), PhLiPS (Pashos et al., 2017), iPSCORE (Panopoulos et al., 2017), CommonMind (Hoffman et al., 2019), Braineac2 (Guelfi et al., 2020), Steinberg_2020 (Steinberg et al., 2021), Young_2019 (Young et al., 2021), Bossini-Castillo_2019 (Bossini-Castillo et al., 2022). It also contains data from the following 7 microarray studies: CEDAR (Momozawa et al., 2018), Fairfax_2012 (Fairfax et al., 2012), Fairfax_2014 (Fairfax et al., 2014), Kasela_2017 (Kasela et al., 2017), Naranbhai_2015 (Naranbhai et al., 2015), Kim-Hellmuth_2017 (Kim-Hellmuth et al., 2017) and Gilchrist_2021 (Gilchrist et al., 2022).

#### Quantification

We quantified transcription at five different levels: (1) gene expression, (2) exon expression, (3) transcript usage, (4) transcriptional event usage, and (5) splice-junction usage (Supplementary Figure 1). Quantification was performed using version v22.05.1 of the <u>eQTL-Catalogue/rnaseq</u> workflow implemented in Nextflow (Di Tommaso et al., 2017). Before quantification, we used Trim Galore v0.5.0 to remove sequencing adapters from the fastq files.

For gene expression quantification, we used HISAT2 v2.2.1 (Kim et al., 2019) to align reads to the GRCh38 reference genome (Homo_sapiens.GRCh38.dna.primary_assembly.fa file downloaded from Ensembl). We counted the number of reads overlapping the genes in the GENCODE V39 (Harrow et al., 2012) reference transcriptome annotations with featureCounts v1.6.4 (Liao et al., 2014). To quantify exon expression, we first created an exon annotation file (GFF) using GENCODE V39 reference transcriptome annotations and dexseq_prepare_annotation.py script from the DEXSeq (Anders et al., 2012) package. We then used the aligned RNA-seq BAM files from the gene expression quantification and featureCounts with flags ‘-p -t exonic_part -s ${direction} -f -O’ to count the number of reads overlapping each exon.

We quantified transcript and event expression with Salmon v1.8.0 (Patro et al., 2017). For transcript quantification, we used the GENCODE V39 (GRCh38.p13) reference transcript sequences (fasta) file to build the Salmon index. For transcriptional event usage, we downloaded pre-computed txrevise (Alasoo et al., 2019; Vija and Alasoo, 2022) alternative promoter, splicing and alternative 3ʹ end annotations corresponding to Ensembl version 105 from Zenodo (https://doi.org/10.5281/zenodo.6499127) in GFF format. These annotations had been augmented with additional experimentally derived promoter annotations from the FANTOM5 consortium (Abugessaisa et al., 2017; FANTOM Consortium and the RIKEN PMI and CLST et al., 2014). We then used gffread (Pertea and Pertea, 2020) to generate fasta sequences from the event annotations and built Salmon indices for each event set as we did for transcript usage. Finally, we quantified transcript and event expression using salmon quant with ‘--seqBias --useVBOpt --gcBias --libType’ flags. All expression matrices were merged using csvtk v0.17.0. Our reference transcriptome annotations are available from Zenodo (https://doi.org/10.5281/zenodo.4715946).

For Leafcutter analysis, splice junctions of the aligned reads were extracted using the *junctions extract* command of the regtools v0.5.2 (Cotto et al., 2023) with options ‘-s $strand -a 8 -m 50 -M 500000’. Then, these splice-junctions were clustered using leafcutter_cluster_regtools.py script from LeafCutter v0.2.9 with options ‘-m 50 -o leafcutter -l 500000 --checkchrom=True’.

#### Normalisation

We normalised the gene and exon-level read counts using the conditional quantile normalisation (cqn) R package v1.30.0 (Hansen et al., 2012) with gene or exon GC nucleotide content as a covariate. We downloaded the gene GC content estimates from Ensembl biomaRt and calculated the exon-level GC content using bedtools v2.19.0 (Quinlan and Hall, 2010). We also excluded lowly expressed genes, where 95 per cent of the samples within a dataset had transcripts per million (TPM)-normalised expression less than 1. To calculate transcript and transcriptional event usage values, we obtained the TPM normalised transcript (event) expression estimates from Salmon. We then divided those transcript (event) expression estimates by the total expression of all transcripts (events) from the same gene (event group). Subsequently, we used the inverse normal transformation to standardise all five molecular quantification estimates. Normalisation scripts together with containerised software are publicly available at https://github.com/eQTL-Catalogue/qcnorm.

### Association testing and statistical fine mapping

We performed association testing separately in each dataset and used a +/− 1 megabase *cis* window centred around the start of each gene. First, we excluded molecular traits with less than five genetic variants in their *cis* window, as these were likely to reside in regions with low genotyping coverage. We also excluded molecular traits with zero variance across all samples and calculated phenotype principal components using the prcomp R stats package (center = true, scale = true). We calculated genotype principal components using plink2 v1.90b3.35. We used the first six genotype and molecular trait principal components as covariates in QTL mapping. We calculated nominal eQTL summary statistics using the GTEx v6p version of the FastQTL (Ongen et al., 2016) software (https://github.com/francois-a/fastqtl) that also estimates standard errors of the effect sizes. We used the ‘--window 1000000 --nominal 1’ flags to find all associations in 1 Mb *cis* window. For permutation analysis, we used QTLtools v1.3.1 (Delaneau et al., 2017) with ‘--window 1000000 --permute 1000 --grp-best’ flags to calculate empirical p-values based on 1000 permutations. The ‘--grp-best’ option ensured that the permutations were performed across all molecular traits within the same ‘group’ (e.g. multiple probes per gene in microarray data or multiple transcripts or exons per gene in the exon-level and transcript-level analysis) and the empirical p-value was calculated at the group level.

We performed QTL fine mapping using the Sum of Single Effects Model (SuSiE) (Wang et al., 2020) implemented in the susieR v0.11.92 R package. We converted the genotypes from VCF format to a tabix-indexed dosage matrix with bcftools v1.10.2. We imported the genotype dosage matrix into R using the Rsamtools v2.8.0 R package. We used the same normalised molecular trait matrix used for QTL mapping. We regressed out the first six phenotype and genotype PCs separately from the phenotype and genotype matrices. We performed fine mapping with the following parameters: L = 10, estimate_residual_variance = TRUE, estimate_prior_variance = TRUE, scaled_prior_variance = 0.1, compute_univariate_zscore = TRUE, min_abs_corr = 0. The steps described above are implemented in the <u>eQTL-Catalogue/qtlmap</u> v22.04.01 Nextflow workflow available from GitHub.

### Filtering of transcript-level summary statistics

We filtered transcript-level summary statistics using a connected components approach (Kolberg et al., 2020) to select the strongest signals per transcript-level group (gene for transcript and exon level, clusters for leafcutter). For each group, first, we filtered out the credible sets where maximum absolute z value is lower than 3 and size is bigger than 200 variants. Then, we found overlapping variants between credible sets, defining these credible sets as connected components. For each connected component we selected the molecular trait with the highest posterior inclusion probability (PIP) and kept only the summary statistics of these selected molecular traits. This approach enabled easier sharing of most significant signals per molecular trait group, decreasing the volume of shared data by 98%.

### Colocalisation with vitamin D GWAS

We used coloc.susie (Wallace, 2021) to perform signal-level colocalisation between all RNA-seq-based datasets in the eQTL Catalogue and GWAS summary statistics for vitamin D levels in the UK Biobank. For all molecular QTLs, we used the log Bayes factors (LBFs) exported by our <u>eQTL-Catalogue/qtlmap</u> v22.04.01 workflow. For the vitamin D GWAS, we used published SuSiE fine mapping results from a previous study (Kanai et al., 2021) downloaded from Google Cloud (<u>link</u>). We performed colocalisation between all pairs of independent fine mapped signals (up to 10 per locus) and reported results where PP4 > 0.9. The colocalisation workflows is available from GitHub (https://github.com/ralf-tambets/coloc).

### Generation of QTL coverage plots

We used the bamCoverage command from deepTools v3.2.0 (Ramírez et al., 2016) with bin-size option ‘-bs 5’ to generate read-coverage (bigwig) files. We then used extractCoverageData and plotCoverageData commands of wiggleplotr R v1.13.1 package (Alasoo, 2017) to read specific regions of the bigwig files, scale all introns to the length of 50 nucleotides, and generate the plots as ggplot2 (Wickham, 2016) objects. Finally, we generated exon QTL effect-size plots with ggplot2 v3.3.6 and put all the plots together with the cowplot v1.1.1 R package (Wilke, 2019). We used *tabix.read.table* from seqminer v8.4 package (Zhan and Liu, 2015) to extract both genotype and QTL data from indexed files in the regions of interest. Coverage plot generation workflow is publicly available at https://github.com/kerimoff/leafcutter_plot.

## Author contributions

N.K. and H.J.T performed quality control of individual-level data and executed the eQTL analysis workflows. N.K. generated the static QTL coverage plots for all molecular QTLs. R.T. modified the genotype imputation workflow to support imputation X chromosome genotypes. P.K. implemented the low-coverage WGS genotype imputation workflow. M.K., and J.C.U performed the fine-mapping analysis on vitamin D levels in the UK Biobank. R.T. implemented the fine mapping analysis for the eQTL Catalogue datasets and performed colocalisation between all molecular traits and vitamin D GWAS. A.V. updated the promoter annotations for txrevise. M.R., J.H., S.G., and D.T. prepared the Braineac2 dataset for eQTL analysis. H.J.T. performed quality control of newly added eQTL datasets. A.C., W.R. and S.K-H. used the eQTL Catalogue workflows to perform federated eQTL analysis on the Kim-Hellmuth_2017 dataset. J.D.H. implemented the eQTL Catalogue summary statistics REST API. I.K., U.R. and H. Peterson set up the eQTL Catalogue credible set browser. I.R. and K.A. performed literature review to prioritise causal genes at fine mapped GWAS loci. K.A, H.Parkinson, A.M and H.F. supervised the project. N.K. and K.A. wrote the manuscript with input from all authors.

## Acknowledgements

The RNA-seq quantification and QTL analyses were performed at the High Performance Computing Center, University of Tartu. We thank O.E. Oopkaup, S. Kuusemets and the rest of the team of the High Performance Computing Center for their professional and timely technical support in enabling the analyses performed in this paper. We thank M. Weale for preparing the Braineac2 dataset for analysis. N.K., J.D.H., P.K. and H.J.T. were supported by a grant from Open Targets (grant no. OTAR2077). H. Parkinson, A.M. and J.H. were supported by the European Molecular Biology Laboratory. K.A., A.C., W.R. and N.K. also received funding from the European Union’s Horizon 2020 research and innovation program (grant no. 825775). K.A., N.K, R.T., A.V., P.K. and I.R. were supported by the Estonian Research Council (grant nos. IUT34-4 and PSG415). K.A. and N.K. were also supported by the Estonian Centre of Excellence in ICT Research (EXCITE), funded by the European Regional Development Fund. I.K., U.R. and H. Peterson were supported by a Distributed Infrastructure for Life-Science Information ELIXIR, European Regional Development Fund project (2014-2020.4.01.16-0271). S.K.-H. was supported by the Emmy Noether Programme KI 2091/2-1 (459153572), SFB/TRR237-B29 (369799452) and SFB/TRR359-B06 (491676693) of the Deutsche Forschungsgemeinschaft (DFG). Funding information for individual studies included in the eQTL Catalogue is presented in Supplementary Note 2.

## Data Availability

The molecular QTL summary statistics, fine mapping results (including SuSiE log Bayes factor) and QTL coverage plots are available from the eQTL Catalogue FTP server (see https://www.ebi.ac.uk/eqtl/Data_access/). The marginal eQTL summary statistics are also available via our REST API (https://wwwdev.ebi.ac.uk/eqtl/api/docs), which we have completely re-written for release 6. RNA-seq and genotype data from the CAP (phs000481.v3.p2), Peng_2018 (phs001586.v1.p1), PhLiPS (phs001341.v1.p1) and iPSCORE (phs000924.v4.p1) studies were downloaded from dbGaP; Steinberg_2020 (EGAD00001005215, EGAD00001003355, EGAD00010001746), Young_2019 (EGAD00001005736) and Bossini-Castillo_2019 (EGAD00001004830, EGAD00010001848) from EGA, and CommonMind (syn2759792) from Synapse. Raw genotype data for Gilchrist_2021 (EGAD00010000144, EGAD00010000520) was downloaded from EGA. Raw gene expression data from Gilchrist_2021 was downloaded from Zenodo (https://doi.org/10.5281/zenodo.6352656). Raw RNA-seq and genotype data from Braineac2 were not deposited.

## Supplementary Note

Reference mapping bias is known to induce false positive associations in splicing and allele specific expression analysis (Kumasaka et al., 2016; Li et al., 2018; van de Geijn et al., 2015). A tool often used to correct for reference mapping bias is WASP (van de Geijn et al., 2015), which has also been included in the STAR (Dobin et al., 2013) RNA-seq short read aligner. Although the eQTL Catalogue uses HISAT2 (Kim et al., 2019) to perform RNA-seq read alignment, we did consider the option to switch to STAR to use WASP read filtering. However, after initial benchmarks we opted against it. First, we found that WASP was very conservative and filtered out a large proportion of reads form exonic regions. As a result, many well-known true positive splicing QTLs were no longer detected and the QTL read coverage plots became noisy due to the large number of filtered reads. Secondly, as implemented in STAR, WASP was only able to account for single nucleotide variants and did not consider short insertions or deletions that have even large potential to cause reference mapping bias. Finally, our transcript usage and txrevise quantification uses Salmon (Patro et al., 2017) to pseudoalign reads directly to the transcriptome and is thus not compatible with WASP. For example, two of the three suspected reference mapping bias cases (*DHCR7* and *NUDT9*) were detected in Salmon transcript usage analysis. Finally, switching to STAR+WASP would have significantly increased the runtime of our RNA-seq quantification workflow which already took over two months at the University of Tartu High Performance Computing Center. For these reasons we decided against directly correcting for reference mapping bias in the QTL mapping process. Instead, we opted to provide access to pre-generated QTL coverage plots that can be used to visually detect strong cases of reference mapping bias.

## Supplementary Note 2

Funding statements for the new studies included in the eQTL Catalogue.

### CommonMind

Bio-samples and/or data for this publication were obtained from NIMH Repository & Genomics Resource, a centralized national biorepository for genetic studies of psychiatric disorders. Data were generated as part of the CommonMind Consortium supported by funding from Takeda Pharmaceuticals Company Limited, F. Hoffman-La Roche Ltd and NIH grants R01MH085542, R01MH093725, P50MH066392, P50MH080405, R01MH097276, RO1-MH-075916, P50M096891, P50MH084053S1, R37MH057881, AG02219, AG05138, MH06692, R01MH110921, R01MH109677, R01MH109897, U01MH103392, and contract HHSN271201300031C through IRP NIMH. Brain tissue for the study was obtained from the following brain bank collections: the Mount Sinai NIH Brain and Tissue Repository, the University of Pennsylvania Alzheimer’s Disease Core Center, the University of Pittsburgh NeuroBioBank and Brain and Tissue Repositories, and the NIMH Human Brain Collection Core. CMC Leadership: Panos Roussos, Joseph Buxbaum, Andrew Chess, Schahram Akbarian, Vahram Haroutunian (Icahn School of Medicine at Mount Sinai), Bernie Devlin, David Lewis (University of Pittsburgh), Raquel Gur, Chang-Gyu Hahn (University of Pennsylvania), Enrico Domenici (University of Trento), Mette A. Peters, Solveig Sieberts (Sage Bionetworks), Thomas Lehner, Stefano Marenco, Barbara K. Lipska (NIMH).

### CAP

The dataset used for the analyses described in this manuscript was obtained from the Cholesterol and Pharmacogenetics (CAP) study through dbGAP (phs000481.v3.p2). Funding support for the generation of this dataset was provided by National Heart, Lung, Blood Institute (NHLBI) grant U01 HL69757. The manuscript was not prepared in collaboration with CAP investigators and does not necessarily reflect the opinions or views of CAP investigators or NHLBI.

### Peng_2018

This work was supported by the National Institutes of Health [NIH-NIMH R01MH094609, NIH-NIEHS R01ES022223, NIH-NIEHS PO1ES022832, NIH-NIEHS R24ES028507, NIH-NIEHS R21ES028226, and NIH-NIEHS R01ES025145. A complete description of the cohort can be found in: Appleton AA, Murphy MA, Koestler DC, Lesseur C, Paquette AG, Padbury JF, Lester BM, and Marsit CJ. Prenatal Programming of Infant Neurobehavior in a Healthy Population. Paediatr Perinat Epidemiol 2016, 30(4): 367-75.

### PhLiPS

This work was supported by grant 5U01HG006398.

### iPSCORE

This work was supported in part by a California Institute for Regenerative Medicine (CIRM) grant GC1R-06673 and NIH grants EY021237, HG008118, HL107442, DK105541 and DK112155. iPSC RNA-seq was performed at the UCSD IGM Genomics Center with support from NIH grant P30 CA023100.

### Bossini-Castillo_2019

This research was funded by the Wellcome Trust (grant number WT206194). L.B.-C. was supported by the MRC Skills Development Fellowship (MR/N014995/1).

### Steinberg_2020

This work was funded by the Wellcome Trust (206194). M.J.C. was funded through a Medical Research Council Centre for Integrated Research into Musculoskeletal Ageing grant (148985). R.A.B. and the Human Research Tissue Bank are supported by the NIHR Cambridge Biomedical Research Centre. J.H.D.B. and G.R.W. are funded by a Wellcome Trust Strategic Award (101123), a Wellcome Trust Joint Investigator Award (110140 and 110141) and a European Commission Horizon 2020 Grant (666869, THYRAGE). A.W.M. receives funding from Versus Arthritis; Tissue Engineering and Regenerative Therapies Centre (21156).

### Young_2019

R.F. was supported by funding from the UK Multiple Sclerosis Society (MS50), the Adelson Medical Research Foundation and a core support grant from the Wellcome Trust and MRC to the Wellcome Trust-Medical Research Council Cambridge Stem Cell Institute (203151/Z/16/Z). A.Y. is supported by a Wellcome Trust Clinicians PhD Fellowship (RRZD/029). All data for this study were generated under Open targets project OTAR039. N.K. and D.J.G. were funded by the Wellcome Trust grant WT206194.

### Gilchrist_2021

C.K. was supported by Wellcome Trust Investigator Award [204969/Z/16/Z], NIHR Oxford Biomedical Research Centre and Chinese Academy of Medical Sciences (CAMS) Innovation 537 Fund for Medical Science (grant number: 2018-I2M-2-002), Wellcome Trust Grants 090532/Z/09/Z and 203141/Z/16/Z to core facilities Wellcome Centre for Human Genetics, Oxford Biomedical Research Computing (BMRC) facility, a joint development between the Wellcome Centre for Human Genetics and the Big Data Institute supported by Health Data Research UK and the NIHR Oxford Biomedical Research Centre. The study was funded by Wellcome Trust Intermediate Clinical Fellowship to B.P.F. (no. 201488/Z/16/Z). J.J.G. is funded by a National Institute for Health Research (NIHR) Clinical Lectureship.

### Braineac2

Mina Ryten, David Zhang, and Karishma D’Sa were supported by the UK Medical Research Council (MRC) through the award of Tenure-track Clinician Scientist Fellowship to Mina Ryten (MR/N008324/1). Sebastian Guelfi was supported by Alzheimer’s Research UK through the award of a PhD Fellowship (ARUK-PhD2014-16). Regina Reynolds was supported through the award of a Leonard Wolfson Doctoral Training Fellowship in Neurodegeneration. All RNA sequencing data performed as part of this study were generated by the commercial company AROS Applied Biotechnology A/S (Denmark).

## Supplementary figures

**Supplementary Figure 1.**
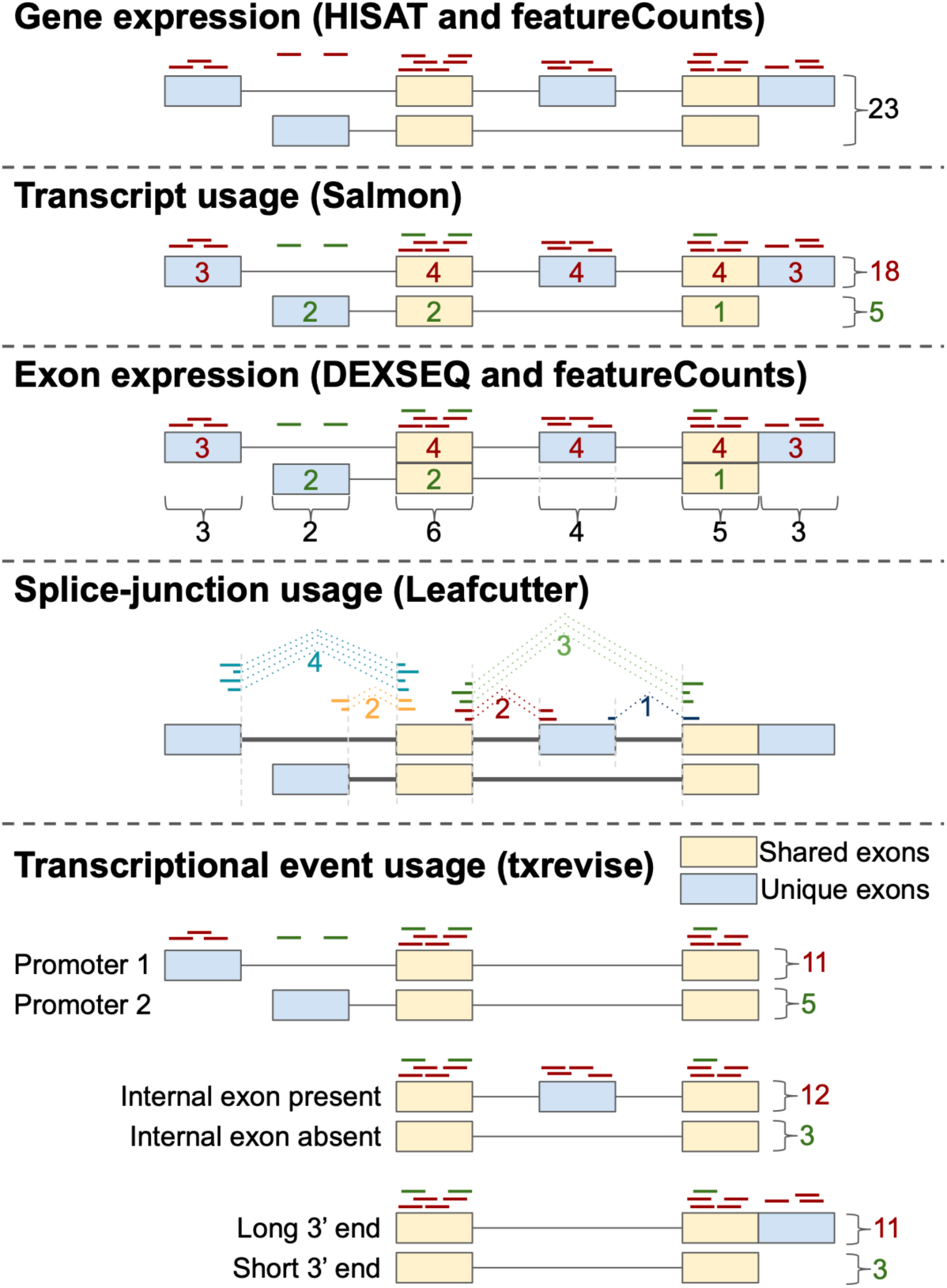
Overview of the five molecular trait quantification methods used by the eQTL Catalogue. Gene expression was quantified by counting the total number of reads overlapping annotated exons of the gene. Transcript usage was estimated with Salmon. Exon expression was estimated by counting the number of reads overlapping each exon. Splice-junction usage was quantified with Leafcutter. Txrevise was used to estimate the expression levels of three types of transcriptional events (promoter usage, splicing and 3′ end usage).

**Supplementary Figure 2.**
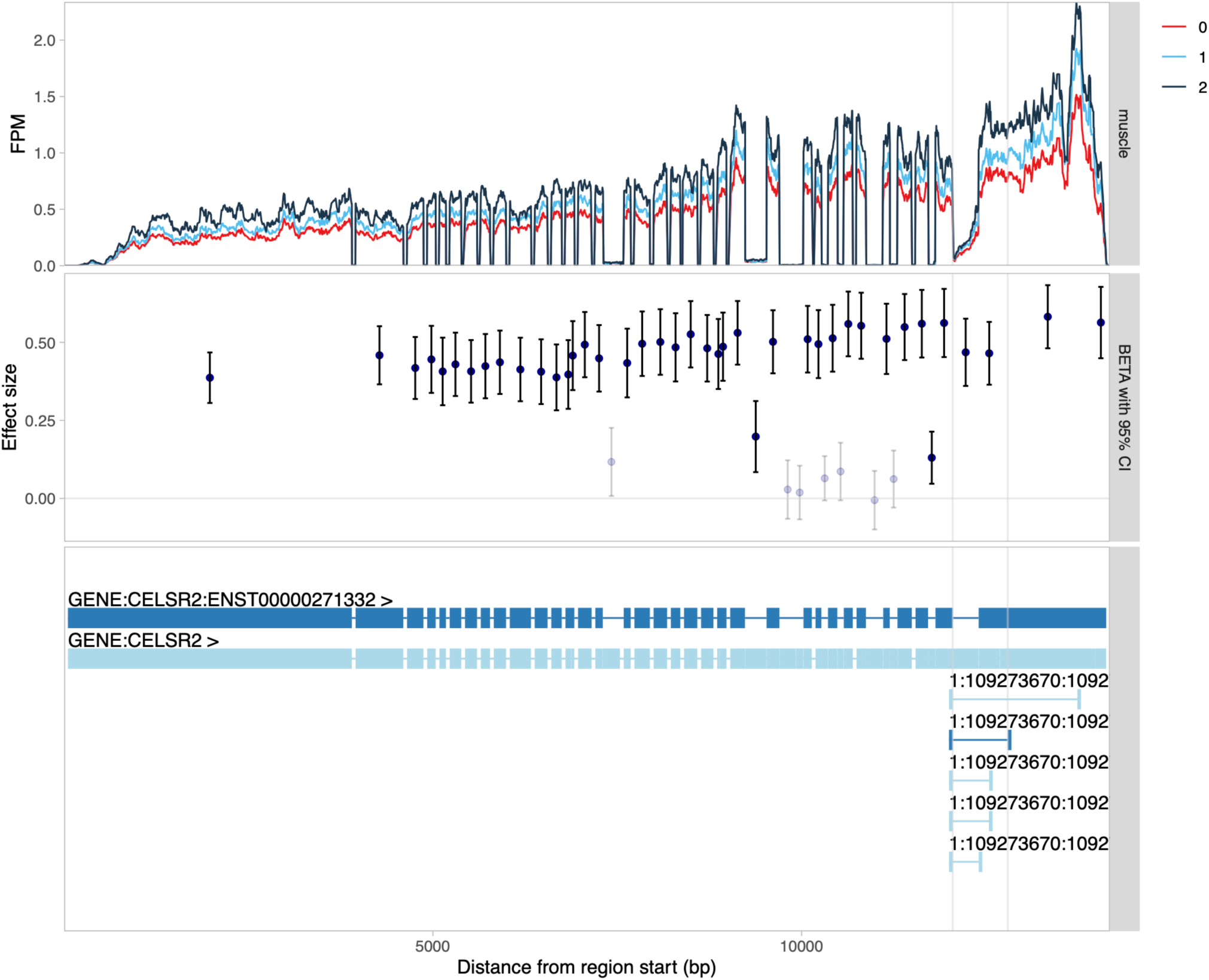
QTL coverage plot for *CELSR2* gene stratified by the lead Leafcutter QTL variant (chr1_109274241_T_TC) in the GTEx muscle tissue. The observed association at junction reads in the 3’ end of the gene is likely a consequence of the strong eQTL effect at this locus rather than the primary mechanism driving complex trait association.

**Supplementary Figure 3.**
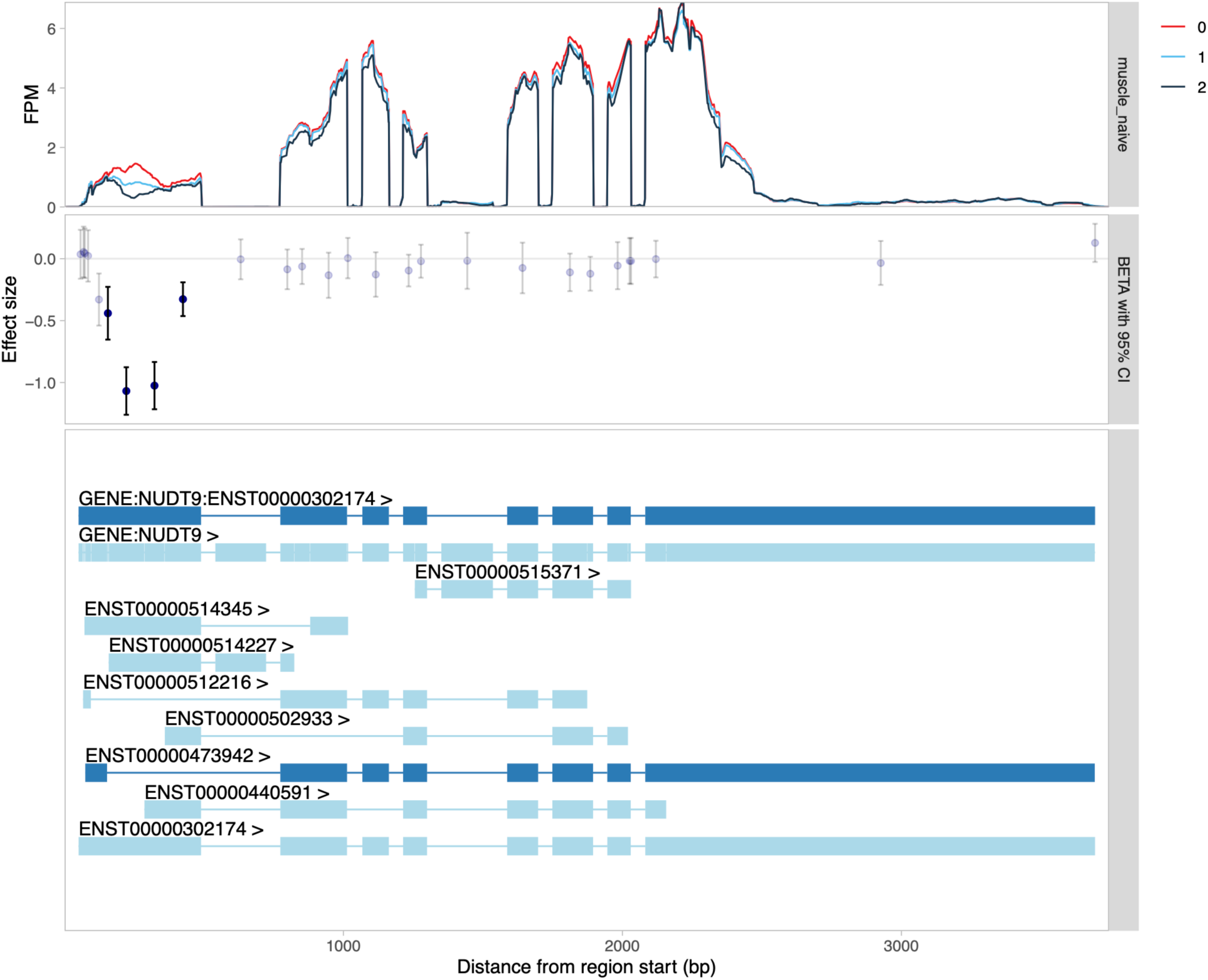
QTL coverage plot for *NUDT9* stratified by the genotype of the lead transcript usage (tx) QTL variant (chr4_87380254_C_T) in the FUSION (Taylor et al., 2019) muscle tissue. The ‘bulge’ in read coverage observed at the 5’ end of the gene suggests potential reference mapping bias.

**Supplementary Figure 4.**
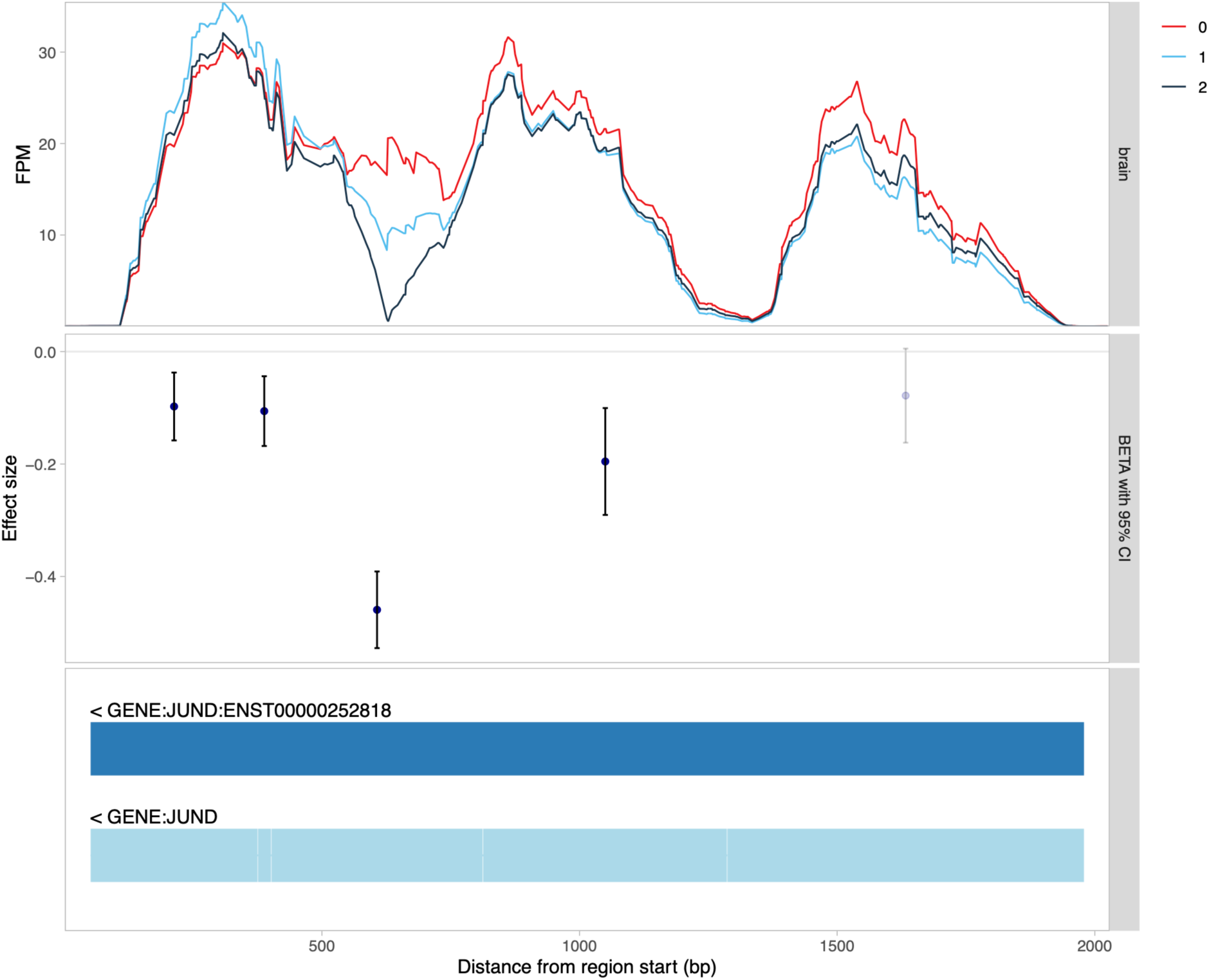
QTL coverage plot for *JUND* stratified by the genotype of the lead gene expression (ge) QTL variant (chr19_18287220_A_C) in the BrainSeq (Jaffe et al., 2018) dataset. The drop in read coverage in the middle of the exon suggests potential reference mapping bias.

**Supplementary Figure 5.**
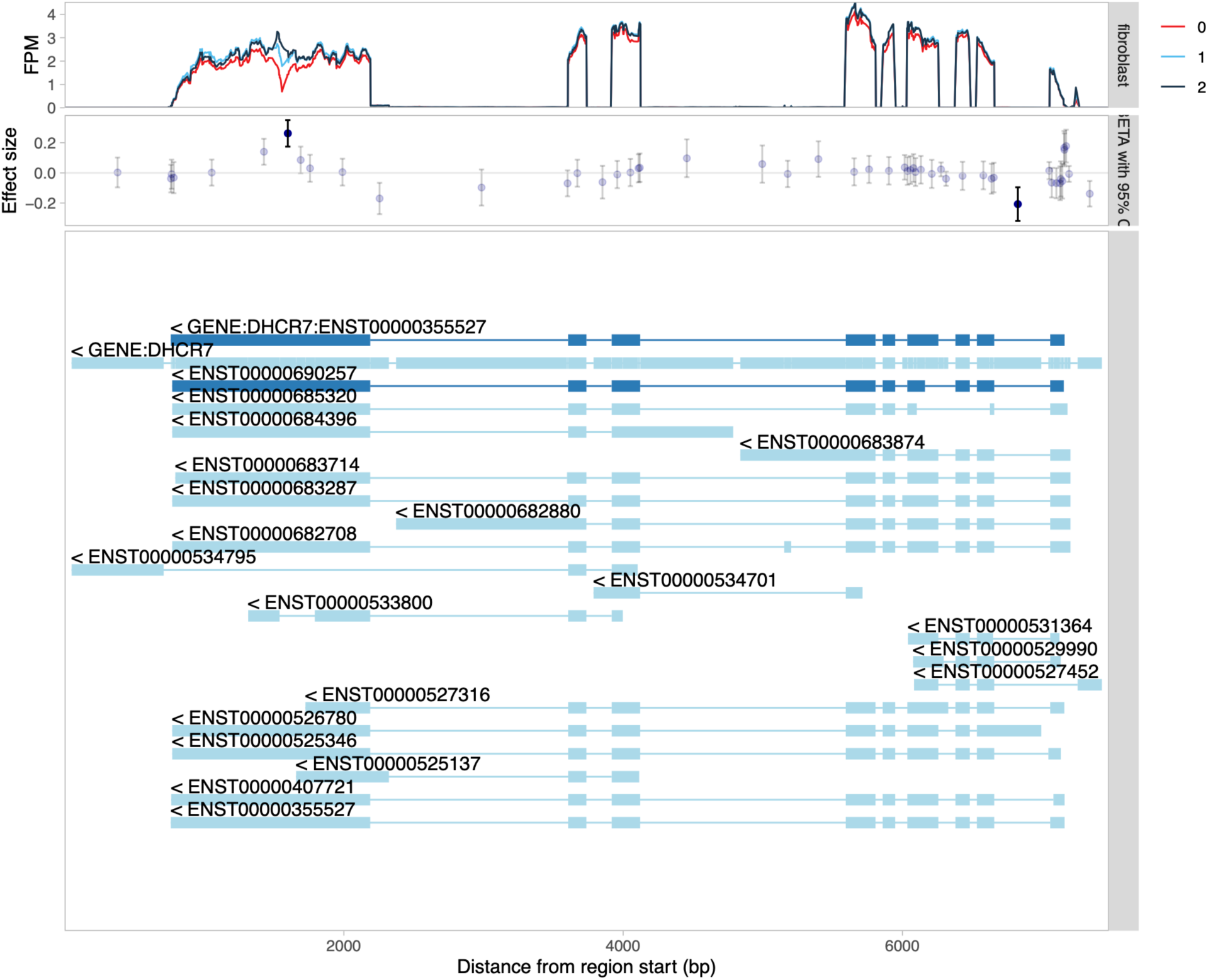
QTL coverage plot for *DHCR7* stratified by the genotype of the lead transcript usage (tx) QTL variant (chr11_71458997_T_C) in the GTEx fibroblast dataset. The ‘bulge’ in read coverage observed at the middle of the last exon of the MANE Select transcript (ENST00000355527) suggests that the association is driven by reference mapping bias.

